# Developmental diversification re-patterns basal antiviral immunity across plant cell types

**DOI:** 10.64898/2026.05.18.725958

**Authors:** Luis Villar-Martín, Brigen Manikan Ardona, Tamara Jimenez-Góngora, Paloma Alvarez-Franco, Kitija Ulme, Victor Manuel González-Miguel, Ignacio Rubio-Somoza

## Abstract

Diversification of plant development has largely enabled land colonization and establishment of different ecosystems. This diversification relies on the gain/loss of cell-types and tissues, such as stomata, vascular tissue and functional roots. Likewise, diversification of immunity is thought to rely on expansion/contraction, followed by functional specification, of the different components of plant defense mechanisms. Although anatomical changes might result in altering the infection routes of pathogens and the cells and tissues they interact with, very little is known about the co-evolution of plant development and immunity. We have recently described that RNAi-dependent antiviral responses observed in the non-*vascular Marchantia polymorpha* are confined to leaf vasculature in *Nicotiana benthamiana* plants, suggesting repatterning of antiviral responses as result of the acquisition of developmental innovations. Here, we explored the genetic basis of that repatterning by establishing the basal immunity toolkit across different cell-types in the non-vascular *Marchantia polymorpha* and the vascular *Arabidopsis thaliana*. The results from our comparative transcriptomic studies show that while RNAi is the major antiviral defense across Marchantia cell-types, that configuration is only maintained in phloem companion cells in Arabidopsis leaves, suggesting that plant immunity might co-evolve with developmental diversification. Additionally, differential levels of RNAi expression in different cell-types correlate with their vulnerability to viral countermeasures, with companion cells been the most resilient to the presence of viral silencing suppressors.

## Introduction

Transition from unicellular to multicellularity, followed by land colonization and the concurrent acquisition of developmental innovations are arguably two of the main steps in plant evolution. Both events majorly contributed to functional specialization and the appearance of new cell identities that led to the emergence of differentiated root and vascular systems, among other morphological novelties. Those developmental innovations not only allowed plants to further spread creating new terrestrial ecosystems but became central for the interaction with microbes that were already inhabiting the surface of our planet, such as the prototypical case of vasculature in plant-virus interactions^1,2^. To fend off potential threats, different plant lineages, as the non-vascular and rootless bryophyte *Marchantia polymorpha* and the vascular *Arabidopsis thaliana*, present conserved immune responses against virus ^3^, bacteria ^4^, oomycetes ^5^ and fungi ^6^. Those conserved responses rely on a plant immune system encompassing three interconnected surveillance systems that monitor the presence of microbe-derived molecules ^7,8^. A highly specialized array of pattern-recognition receptors (PRRs) perceives pathogen-associated molecular patterns (PAMPs), such as bacterial flagellin, fungal chitin and viral double-strand RNA (dsRNA) at the extracellular space, activating pattern-triggered immunity (PTI) ^9,10^. PRR proteins can be classified according to their structural features, encompassing a diverse set of ectodomains, such as leucine rich repeats (LRRs), lectins or lysin motifs (LySM), and the presence of a cytosolic kinase domain in receptor-like kinases (RLKs) or its absence in receptor-like proteins (RLPs). Both RLKs and RLPs share a common evolutionary origin^11^. Additionally, plant cells bear two intracellular surveillance systems that sense foreign molecules and/or their effect on disturbing cellular homeostasis. While nucleotide-binding leucine-rich repeat (NLR) receptors monitor foreign proteins, the RNA silencing machinery tracks double-stranded RNAs (dsRNA) within host cells. NLR proteins are classified based on the presence of different protein domains in their N-terminal region and enact on activation effector triggered immunity (ETI)^12^. The presence of dsRNA is recognized by members of the Dicer-like family (DCL) triggering the so-called RNA interference (RNAi) immunity. DCLs process dsRNA into small RNA duplexes typically ranging from 20 to 25 nucleotides in length. sRNA duplexes are subsequently loaded into the RNA silencing complex, where proteins from the ARGONAUTE family are the main executors, that scans the intracellular space for nucleic acids with high sequence complementarity to abrogate their function. dsRNA can be produced as replication intermediates of RNA viruses, products of viral transcription and the action of host RNA-dependent-RNA polymerases (RDRs) that convert single-strand RNA (ssRNA) molecules into dsRNA^13^. To thwart these plant surveillance and defense systems pathogens have evolved molecular countermeasures, generically called effectors, that they deliver into host cells. Concomitantly, plants diversified their immune mechanisms through expansion/contraction and specialization of the elements involved, such as PRRs, NLRs, DCL, RDR and AGO^11,14-18^.

Despite our increasing knowledge of the diversification trajectories of plant development and immunity, the extent of their co-evolution remains poorly understood. Seminal studies have shown that RNAi-mediated responses induced by viral infection in the non-vascular plant Marchantia have been specifically recruited to vascular tissues in Nicotiana species^3^. These results suggest that newly acquired tissues and/or cell types may have driven the rerouting of defense responses by acting as turning into focal points for pathogen interaction. hence contributing to the evolution of plant immunity.

In this study we aimed to dissect the genetic basis underlying the cellular diversification of antiviral responses between non-vascular and vascular plants. Our comparative cell-type transcriptomic approach shows that spatial relocation of those responses is based on the prepatterned expression of their immune elements. While core members of the RNAi machinery exhibit a broad basal expression pattern in *Marchantia polymorpha*, their expression is concentrated in the vascular companion cells in *Arabidopsis thaliana*. Interestingly, RNAi is consistently the main contributor to basal immunity in all Marchantia cell-types whereas that immune configuration is only found in Arabidopsis companion cells, with PRR and NLR-based defense is more abundant in other leaf cell-types. Different levels of overall expression of RNAi components in specific cell-types in Arabidopsis leaves correlate with resilience to the interference with their defense function by viral silencing suppressors, with companion cells displaying the highest resistance to perturbations in RNA silencing.

## Results

### RNAi is the prevailing antiviral system in all Marchantia’s cell-types

We recently showed that antiviral RNAi-mediated responses to TMV infection in the non-vascular plant *Marchantia polymorpha* resemble those exclusively found in the vasculature of *Nicotiana benthamiana* plants^3^. These findings prompted us to investigate the underlying mechanism of this antiviral response diversification. To this end, we revisited the transcriptional profile of mock treated Marchantia plants reported by Ros-Moner *et al*. (2024)^3^ to determine whether the observed response arose from de novo induction or elevated basal expression of core RNAi response genes. We found that both MpDCL4 and MpRDR6 genes are basally expressed in mock treated plants (Figure 1A).

**Fig. 1:**
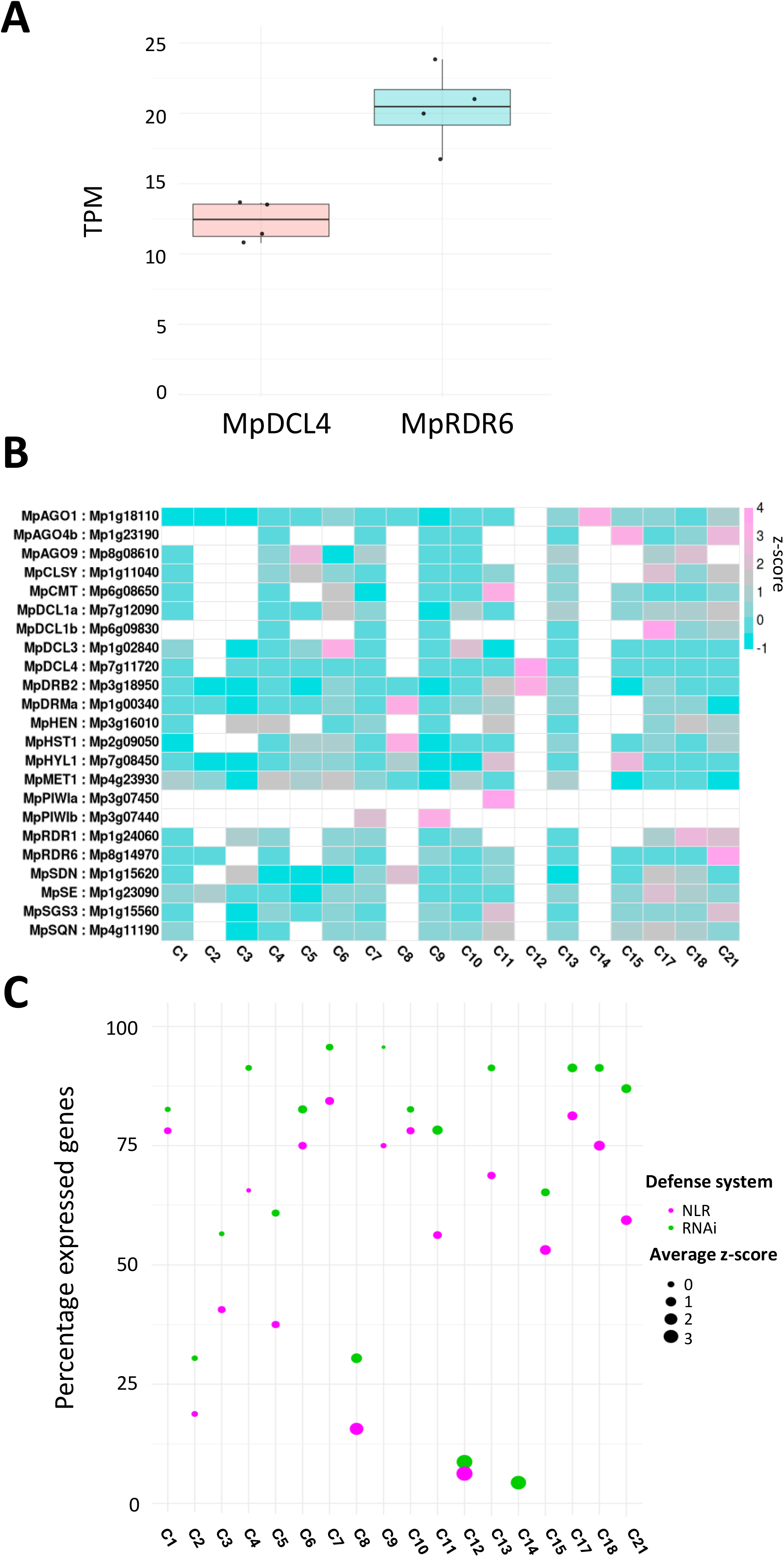
Members of the RNAi machinery have a broad basal expression and are the major contributors to cell-type specific antiviral defense *in Marchantia polymorpha*. **A**. Basal expression levels of two of the core RNAi elements for antiviral defense, MpDCL4 and MpRDR6 in 28 days-old plants. **B**. Heatmap of expression of RNAi members in different cell-types from 31 days-old plants. Only RNAi components showing expression in at least one cell-type are shown. **C**. Relative contribution of RNAi and NLR members to the overall cell-type specific antiviral defense. Y-axis shows the percentage of genes expressed from each pathway, thickness of dots shows the average Z-score expression of the elements from each pathway in each cell type.

To better define the expression pattern of MpRDR6, we reanalyzed the single cell RNA-seq (scRNA-seq) data from 31 days old Marchantia plants generated by Wang *et al*. (2023)^19^. MpRDR6 expression was detected in 13 of the 18 cell clusters identified at this developmental stage, by using a threshold of ≥ 1 transcript per million (TPM; Figure 1B). Altogether, these results indicate that *MpRDR6* is broadly expressed in wild type Marchantia plants.

We additionally reannotated other components of the RNAi pathways in *Marchantia polymorpha* based on their orthology to those in the *Arabidopsis thaliana* genome (Supplementary Table 1) and assessed their expression patterns using the same criteria applied to *MpRDR6*. We found that at least 23 out of the 26 analyzed genes were expressed in at least one cell cluster, with most of them displaying broad expression across all cell populations. Notable exceptions were pattern *MpPIWIa* and *MpPIWIb*, which were detected in only one and two clusters respectively (Figure 1B). These results suggest that RNAi activity is widespread in all Marchantia cell-types.

We then sought to determine how this above-described broad expression pattern of RNAi-related components compares with that of the two other plant antiviral surveillance systems mediated by plasma membrane PRRs and intracellular NLRs. Using the same criteria as above, we found that 32 out of the 41 NLR genes in the Marchantia genome^20^ were expressed in at least one cell cluster (Supplementary Figure 1, Supplementary Table 1). Interestingly, cell cluster 14 lacked any NLR expression. To further assess the relative contribution of RNAi and NLR-based immune systems to the overall antiviral defense across cell populations, we normalized the total number of genes in each system which were expressed in at least one cluster to 100% (23 RNAi and 32 NLR genes), and calculated the proportion expressed in each cluster (Figure 1C). This approach revealed a consistent predominance of the RNAi system across all cell clusters. This trend was largely maintained upon inclusion of PRRs with the only exception of cluster 2 where PRRs are more abundant (Supplementary Figure 2, Supplementary Table1). In total, 129 of the 191 PRRs (125 RLKs and 66 RLPs) encoded in the Marchantia genome^18^ were expressed in at least one cell cluster, including 29 RLPs and 100 RLKs (Supplementary Figure 3).

Altogether, these results indicate that each Marchantia cell type harbors specific antiviral defense repertoires, with RNAi being the most predominant system.

### Prevalence of RNAi mediated antiviral immunity is restricted to companion cells in Arabidopsis leaves

Given the broad cellular distribution of the antiviral RNAi defense machinery in Marchantia plants, we asked whether a similar pattern is observed in vascular plants. To that end, we obtained the transcriptomic profile of three of the most important leaf cell-types for plant-virus interaction in *Arabidopsis thaliana* and viruses, such as mesophyll cells, bundle sheath and companion cells. These cells were specifically isolated using GFP reporter lines for these tissues (Supplementary Figure 4) by coupling protoplast generation followed by fluorescence-activated cell sorting (FACS). Using an expression threshold of ≥ 1TPM in at least two of the three biological replicates, with a mean expression also exceeding 1TPM, we identified 14,562 genes expressed in mesophyll cells, 15,340 in bundle sheath and 15,767 in companion cells. A large proportion of these transcripts (13,150) were shared across all three cell-types (Supplementary Figure 5). Among cell type-specific transcripts, the higher number was observed in companion cells (1,288) followed by those found in bundle sheath (603) and mesophyll cells (432). Notably transcriptome similarity between cell-types reflected their spatial proximity in leaf organs (Supplementary Figure 5). Gene ontology analysis of the cell type-specific genes revealed an enrichment for secondary metabolism pathways (glucosinolates and sulfur) in BS^21^, whereas genes associated with phloem development and vegetative phase transition, including the flowering time regulators FT^22^, FD^23,24^ and SPL5^25,26^ were enriched in CC (Supplementary Figure 5). Analysis of the expression pattern of RNAi-related genes (≥ 1TPM) revealed a clear enrichment in companion cells, where members of the RDR gene family were exclusively expressed (Figure 2A, Supplementary Table 2). RDR6 expression pattern was further confirmed using AtRDR6p:GUS-RDR6 3’UTR reporter lines (Figure 2B), while expression of RDR1 and CLSY1 restricted to the Arabidopsis leaf vasculature has been also previously reported^27,28^. To assess the contribution of the other two basal defense immune systems, we established the expression profile of NLRs and PRR encoding genes^18,29^. Using the same criteria as for RNAi-related genes, we found that 120 of 214 annotated NLRs^29^ were expressed in at least one cell-type (Supplementary Figure 6, Supplementary Table 2). After normalizing the total number of expressed genes per system to 100%, we compared the relative contributions of RNAi and NLR pathways across cell types. NLRs were most prevalent in mesophyll and bundle sheath cells, whereas RNAi was the dominant defense system in companion cells (Figure 2C). This pattern was maintained on inclusion of PRRs in the analysis. In total, 263 (52 RLPs and 211 RLKs) of 473 PRRs (93 RLPs and 380 RLKs)^18^ were expressed in at least one cell-type (Supplementary Figure 7, Supplementary Table 2) with an intermediate contribution respective to NLRs and RNAi (Supplementary Figure 8).

**Fig. 2:**
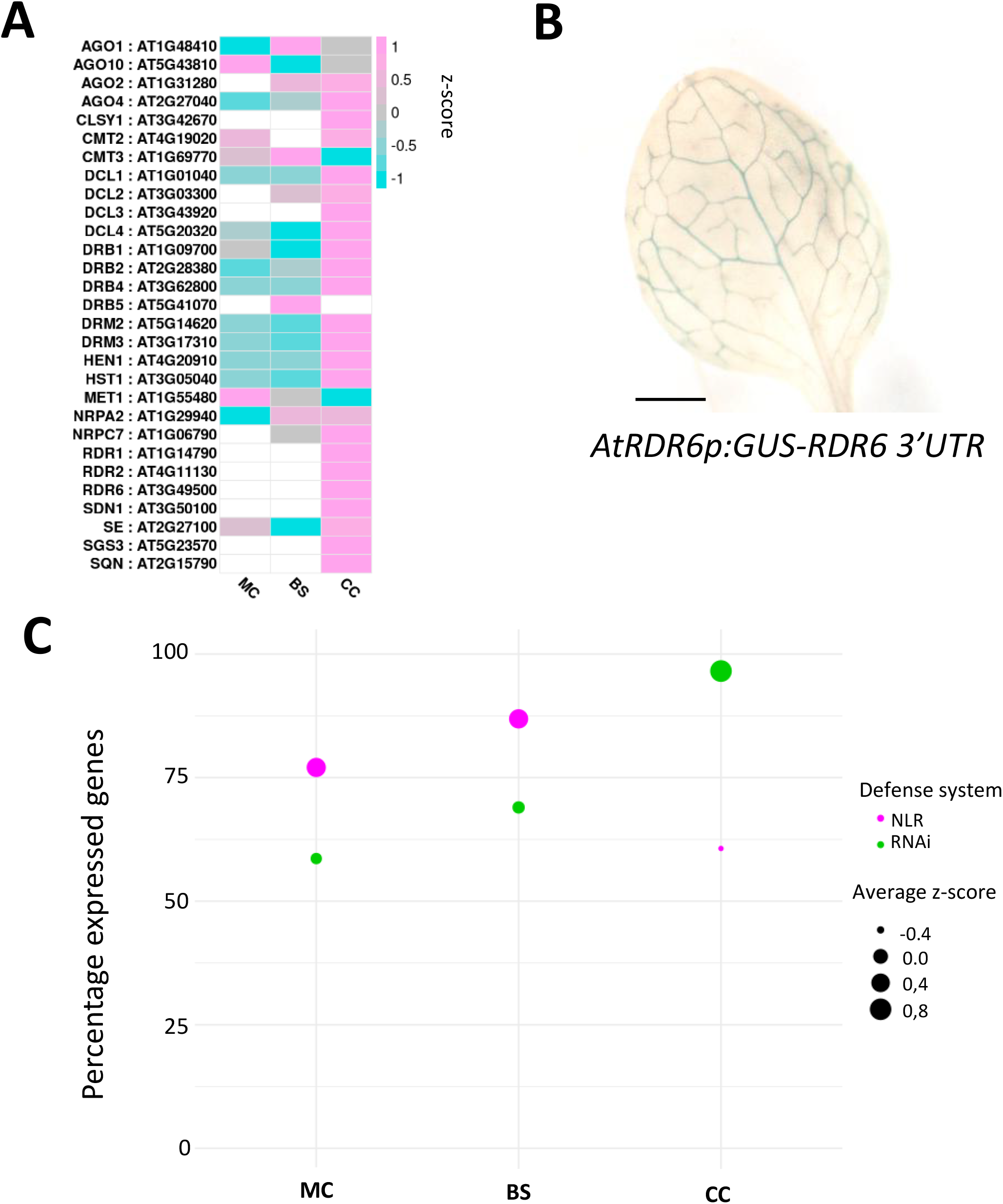
RNAi machinery expression is enriched in phloem cells where is the major contributor to antiviral defense in leaves from *Arabidopsis thaliana*. **A**. Heatmap of expression of RNAi members in different cell-types from Arabidopsis leaves. Only RNAi components showing expression in at least one cell-type are shown. **B**. Transcriptional activity of RDR6 regulatory sequences is confined to the vasculature. **C**. Relative contribution of RNAi and NLR members to the overall cell-type specific antiviral defense. Y-axis shows the percentage of genes expressed from each pathway, thickness of dots shows the average Z-score expression of the elements from each pathway in each cell type. MC=mesophyll cells, BS=Bundle sheath cells and CC= Companion Cells. Scale bars 0.5 cms.

Together, these results show a cell-type specific contribution of the three main antiviral defense mechanisms with RNAi being specifically restricted in companion cells.

### Companion cells are refractory to the viral-derived silencing suppressors

During infection, viruses deploy countermeasures to suppress RNAi-based host defenses. In turn, our results above showed that different plant cell-types exhibit distinct levels of basal RNAi protection, suggesting that cell-types with higher basal RNAi immunity may be more refractory to viral silencing suppressors than those with lower activity. To test this hypothesis, we sought to identify which cell types are targeted for RNAi intervention during infection with the ssRNA turnip mosaic virus (TuMV). We infected to this purpose Arabidopsis lines bearing a reporter system that monitors RNAi integrity, with a TuMV isolate encoding the ROS1 transcription factor, which induces anthocyanin accumulation and thus facilitates visualization of infection^30^. The Arabidopsis reporter system consists of a constitutively expressed artificial miRNA (amiR) targeting a nuclear-localized scarlet (NLS-Scarlet reporter; Supplementary Figure 9). Upon infection, we observed increased nuclear Scarlet signal in mesophyll, bundle sheath and companion cells compared to mock treated controls, hence indicating that these cells are targeted for RNAi dysfunction during systemic viral infection (Figure 3A). We then next challenged the RNA silencing machinery in these cell-types by expressing the TuMV silencing suppressor P1/Hc-Pro, which targets core elements of the RNAi machinery both in vascular and non-vascular plants^31-33^. The percentage of plants displaying phenotypic alterations upon P1/Hc-Pro expression, inversely correlated with the basal RNAi basal activity in those cell-types. Specifically, expression in mesophyll and bundle sheath cells resulted in 37% and 28% of plants, respectively, as compared to only 11% of plants with morphological alterations, when expressed in companion cells (Figure 3B). Cell-type specific induction of P1/Hc-Pro, using the GR-LhG4/pOp system^34^, in Arabidopsis amiR-Scarlet reporter lines confirmed that mesophyll cells are the most sensitive to viral silencing supperssors (Supplementary Figure 10). Next, we analyzed the specific response of these cells by assessing their transcriptional changes after P1/Hc-Pro induction, by combining GFP-assisted cell isolation with transcriptome profiling. To that end, we introduced the GR-LhG4/pOp constructs driving P1/Hc-Pro expression in mesophyll, bundle sheath and companion cells, in the corresponding NLS-GFP marker lines (Supplementary Figure 4). Notably, mesophyll cells exhibited 151 differentially expressed genes (DEG, padj≤0,1) compared to control plants, whereas only 7 DEG were detected in companion cells (Figure 3C, Supplementary Table 3), despite showing comparable levels of expression of P1/Hc-Pro (Figure 3D).

**Fig. 3:**
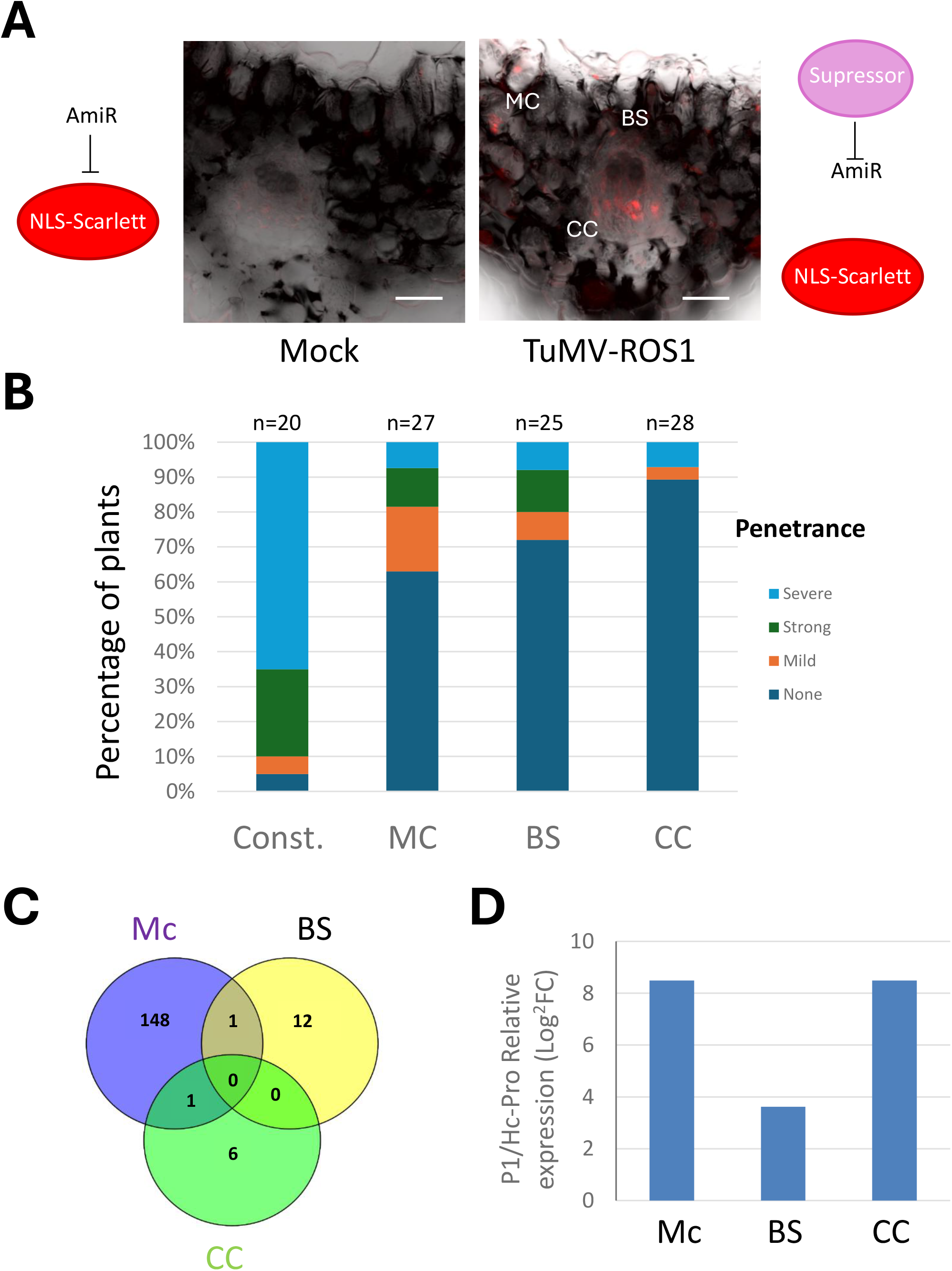
Cell-type specific levels of RNAi machinery expression endorse them with differential reactivity to silencing suppression. **A**. TuMV infection impairs RNAi functioning in cells from Arabidopsis leaves as inferred from higher levels of nuclear scarlet signal (right) when compared to uninfected plants (left). Images show the overlap of bright field and red channels. **B**. Penetrance of developmental phenotypes resulting from P1/Hc-Pro expression constitutively (Const, HRT5 promoter), mesophyll cells (MC, CAB3 promoter), Bundle sheath cells (BS, Sultr2;2 promoter) and companion cells (CC, SUC2 promoter). The number of plants per genotype included in the assay are on top of each column (n=X) **C**. Venn diagram of DEG upon P1/Hc-Pro induction in each cell type. D. Relative induction of P1/Hc-Pro expression levels in the different cell types under study. MC=mesophyll cells, BS=Bundle sheath cells and CC= Companion Cells. White scale bar signifies 40 µMs.

Overall, these results indicate that the impact of viral countermeasures targeting host RNA silencing machinery is cell-type specific and relates to the basal contribution of this antiviral defense mechanism in different cell-types.

## Discussion

Our results reveal a contrasting spatial organization of RNAi-mediated antiviral defense among plant lineages. In the non-vascular plant *Marchantia polymorpha*, components of the RNAi machinery are broadly expressed across somatic cell types. In contrast, in Arabidopsis their basal expression is largely restricted to vascular companion cells. When compared with the basal distribution of the two other major antiviral systems, PPRs and NLRs, RNAi is found to be the predominant defense mechanism across all *Marchantia* cell-types. In Arabidopsis, however, that prevalent configuration is retained only in companion cells and NLRs outweigh the antiviral basal defense contribution of PRRs and RNAi in leaf mesophyll and bundle sheath cells. Importantly, differences in RNAi basal defense distribution correlate with the cell-type capacity to overcome pathogen countermeasures. Thus, mesophyll cells which display the lowest RNAi basal levels, are the most susceptible to viral-derived silencing suppressors, whereas companion cells, with the highest RNAi contribution, are the most refractory, thereby showing that the RNAi machinery spatial configuration influences the outcome of host-virus interaction at the cellular level.

The diversification of plant developmental trajectories involved the acquisition and loss of tissues together with the specialization of cell-types, enabling evolutionary shifts in life strategies such as auto- and heterotrophy. Recent technological advances allowed the identification of previously unknown plant cell types, resolve the heterogeneity of known cell-types and reconstruct the incremental evolutionary complexity of plant tissues through the discovery of foundational genes^35^. In stark contrast, how these developmental innovations relate to changes in plant immune strategies remains poorly understood. Core defense mechanisms against viruses, bacteria, fungi, and oomycetes are broadly conserved across plant lineages^3-6^, recent studies also showing specific defense strategies^36,37^. Increasing evidence also points that different plant cell-types, and even specific cells, deploy characteristic defense responses against pathogens^29,38,39^. The limited data available suggests that basal immune configurations have been shaped by the likelihood of pathogen encounter, to cope with pathogen threats. Thus, PRR receptors recognizing bacterial flagellin and polyglucans, as well as those perceiving fungal chitin, are preferentially expressed in Arabidopsis and Marchantia in cell-types more exposed to microbial infection^40,41^.

We have recently shown that RNAi-dependent antiviral defense responses in the non-vascular bryophyte *Marchantia polymorpha* resemble those restricted to vascular tissues in *Nicotiana benthamiana* leaves, suggesting that acquisition of developmental innovations, that become focal to the interaction with pathogens, has driven the anatomical repatterning of immune responses^3^. Our current results extend this concept by showing that such repatterning not only occurs at the response level, but also at the basal level of expression of core RNAi machinery elements. While RNAi components are broadly distributed across somatic cell-types in adult Marchantia plants, their expression is spatially confined to vascular companion cells in Arabidopsis leaves. This indicates that beyond expansion and contraction of the gene families encoding the three main defense mechanisms, and their functional specialization, plant immunity has also diversified through changes in the spatial pattern of distribution of its core components. Consistent with this view, the relative contribution of RNAi, PRR, and NLR pathways to basal defense differs across each cell type between plant lineages. Thus, while *Marchantia* cell-types predominantly rely on RNAi as their main basal defense system, *Arabidopsis thaliana* exhibits a more dynamic cell-type configuration in which RNAi dominance is restricted to companion cells. We also found that the cell-type specific contribution of RNAi correlates with resilience to viral countermeasures, reinforcing the idea that basal defense patterning and configuration is shaped by the selective pressure imposed by pathogen exposure and tissue-specific interaction, such as the acquisition of the vascular system as a focal site for virus invasion and the enrichment of RNAi in that tissue.

At the same time, restricting RNAi immunity to specific cell-types, as observed in Arabidopsis leaves, may come at the cost of increased vulnerability to pathogen-imposed perturbations of other cell-types. Thereby, the biological reason for reducing the levels of RNAi basal expression in mesophyll cells, deserves further research.

Finally, our work provides detailed cell type-resolved atlases of the immune components in two plant species with contrasting anatomy and living strategies, offering a valuable resource to the identification of novel defense mechanisms and gaining a deeper understanding of how immune functions are partitioned across cells confronting different pathogens. Thus, those atlases would enable rational intervention within the immune toolkits of those specific cell-types a pathogen is interacting with in the course of infection.

## Methods

### Plant growth conditions

*Arabidopsis thaliana* plants were grown either on half strength Murashige Skoog medium (Duchefa) containing 0.8% agar or in soil under long day conditions at 23ºC.

### Constructs and plant transformation

Genomic DNA from *Arabidopsis thaliana* plants was used for amplifying 1997 pbs upstream of the start codon and 545 pbs after the stop codon for the 3’UTR region of AtRDR6 (At1g49500) using primers listed in Supplementary Table 4 to generate the AtRDR6 transcriptional reporter construct using the Green Gate technology^42^. Cell-type marker lines were also generated using the Green Gate technology and cloning a nuclear localization signal, translationally fused to GFP, or tdTomato, under the control of the CAB3p^43^ (At1g29910, for mesophyll cells, 1556 pbs upstream of the start codon), Sultr2;2p^44^ (At1g77990, for bundle sheath cells, 3385 pbs upstream of the start codon) and SUC2p^45^ (At1g22710, for companion cells, 2222 pbs upstream of the start codon) from *Arabidopsis thaliana* using primer listed in Supplementary Table 4. CAB3 and Sultr2;2 promoters were domesticated by overlapping PCRs to remove Eco31I restriction sites with primers detailed in Supplementary Table 4.

AmiR-scarlet reporter system was generated by the following the Green Gate “Two expression cassettes on one T-DNA” approach. amiR scarlet was designed using WMD3 (http://wmd3.weigelworld.org/cgi-bin/webapp.cgi) to specifically recognize the scarlet sequence. Both amiR and NLS-scarlet were under the control of the HTR5 (At4g40040) constitutive promoter from Arabidopsis.

P1/Hc-Pro coding sequence was amplified from DNA isolated from plants overexpressing P1/Hc-Pro described in^46^ and cloned using the Green Gate technology into the final vectors under the control of HTR5, CAB3, Sultr2;2 or SUC2 promoters alone, or in combination with the GR-LhG4/pOp inducible system using the same strategy for cloning two expression modules in one T-DNA. Control constructs for inducible assays were created by cloning Scarlet gene into the GR-LhG4/pOp inducible system using the same strategy for cloning two expression modules in one T-DNA and under the control of CAB3, Sultr2;2 or SUC2 promoters.

All constructs used in this study can be found in Supplementary Table 5.

Arabidopsis lines harboring an empty vector, and P1/Hc-Pro overexpressor, were described previously in^46^.

*Arabidopsis thaliana* plants were transformed following^47^.

### Plant infections

Arabidopsis plants carrying the amiR-scarlet reporter system were grown for 7 days under LD conditions. A subset of plants was kept as an uninfected control, while the rest of the plants were infected by piercing their two cotyledons with a hypodermic needle previously submerged in a solution of agrobacterium (0,6 OD in 10 mM MgCl_2_, 10mM MES, 150 µM acetosyringone, pH5.7) bearing a vector containing the TuMV genome including the coding sequence for ROS1^30^. Plants grew further for two weeks, and purple positive leaves were collected from infected plants along with equivalent ones from uninfected control plants.

### Histology and imaging

Arabidopsis plants were fixed in 90% acetone and GUS activity was assayed as described in^48^. Staining reaction was finished by successive dehydration in solutions with increasing ethanol (20%, 35%, 50%) and fixed in FAA for 30 minutes before storage in Ethanol 70%

For visualization of sRNA reporter activity in uninfected and TuMV-ROS1 plants, individual leaves were embedded in 4 % agarose and cross-sections of around 200 µm were obtained using a vibratome. Confocal imaging was performed using a Leica SP5.

### Cell sorting and RNA isolation

Homozygous plants containing the cell-type specific NLS-GFP markers were crossed to homozygous lines carrying the respective scarlet or P1/Hc-Pro inducible constructs selected for not having basal expression and a good induction of both genes, as assayed by Rt-qPCR. The resulting F1 Arabidopsis plants were grown for 10 days under LD conditions and 22º C. After spraying with 10 µM dexamethasone supplemented with 0.02% Silwett L-77, the third leaf from 11- and 14-days old plants were collected. Leaves were cut into 0.5-1mm strips with a clean scalped blade, and the pieces were transferred and submerged into the enzyme solution (0.4 M Mannitol, 20 mM KCl, 20 mM MES buffer pH 5.7, 1.5 % Cellulose R-10, 0.4 % Macerozyme R-10, 0.35 % Pectolyase Y-23, 10 mM CaCl2, 0.1 % BSA). After an incubation of 1 hour and 30 minutes at 80 rpm, protoplasts were released by gentle circular agitation of the plate and the solution was filtered through a 100 or 70 μm filter, depending on the cell-type to be assayed (mesophyll or vascular bundles, respectively). Protoplasts were washed with W5 solution (154 mM NaCl, 125 mM CaCl2, 5 mM KCl, 2 mM MES buffer pH5.7). After a centrifugation for 3 minutes at 1000 rpm at room temperature, protoplasts were washed one more time and resuspended in 500 μl of W5 solution. 2000 GFP-positive protoplasts per replicate were isolated using the MoFlo XDP cell sorter. Protoplasts were collected in Eppendorf tubes containing RLT buffer (QIAGEN RNeasy Micro Kit) supplemented with 1 % of β-mercaptoethanol and immediately stored in dry-ice. Total RNA from protoplasts was isolated using the QIAGEN RNeasy Micro Kit, following manufacturer instructions.

Total RNA quality and concentration were determined by using bioanalyzer RNA Pico chips and Qubit fluorometer.

### RNA sequencing and expression analysis

*Quantification of basal expressions of MpRDR6 and MpDCL4 in RNA-seq mock treated samples from (Ros-Moner et al 2024) was performed as follows. 1 ug of total RNA was used for preparing each stranded library by mRNA enrichment following manufacturer indications (TrueSeq kit from Illumina). Libraries were sequenced on a Novaseq 6000 device. A total of 20–21 million pairs were obtained per sample. Adapter removal and trimming of low quality bases was done using Trim Galore! v0.6.1 (https://www.bioinformatics.babraham.ac.uk/projects/trim_galore/), removing reads shorter than 20 bps and non-paired reads after the trimming step. Ribosomal RNA reads were removed using SortMeRNA v2.1b, removing also reads that become unpaired after this filtering step. Cleaned reads together with the transcriptome of Marchantia polymorpha were used to quantify gene expression at transcript level using Salmon v1.1.0^49^. For this, the file MpTak1v5.1_r1.mrna.fasta containing the Marchantia transcriptome (v5.1) was downloaded from https://marchantia.info/download/tak1v5.1/ (as of 19-Jan-2021). This file was indexed using Salmon index and then used as input for Salmon quant, which was ran with parameters -l A, –validateMappings, --recoverOrphans, --rangeFactorizationBins 4, --seqBias, and –gcBias. Mapping rate was found to be in the 89.5%–92.58% range. The R progam tximport v1.14.2^50^ was used to aggregate Salmon’s transcript expression estimates at gene level. Figure was obtained using ggplot2.*

Members of the RNAi machinery in Marchantia were identified by blasting the protein sequence of their orthologs in Arabidopsis in on-line available tools (https://plantregmap.gao-lab.org/id_mapping.php and https://marchantia.info/tools/blast/marchantia/). For quantification of the expression levels in the different cell-types from Marchantia 31 day old plants the Seurat object was downloaded using ftp://download.big.ac.cn/gsa2/CRA009114/CRR622988/CRR622988.tar.gz

The object contained data for 15,601 features across 46,136 samples. A pseudobulk analysis was done using the *AggregateExpression* function from *Seurat* 5.2.1 loaded in R4.3. In this process, the total gene counts are aggregated across cells. The resulting pseudo object contained data for 20 clusters for all six time conditions. Day 31 corresponded to the sample labelled S0, for which count data was found in clusters 0 to 14, 16, 17 and 20. Then, normalization was applied using Seurat’s N*ormalizeData* function with parameters *normalization*.*method = “RC”* and *scale*.*factor = 1e6* [The feature counts for each cell are divided by the total counts for that cell and miltiplied by the scale factor]. This produces counts-per-million (CPM) values which, being this data from a 3’-scRNAseq library preparation, are equivalent to transcripts-per-million (TPM) values.

The table of bulk TPM data was then exported and cluster labels were modified so that cluster 0 was renamed cluster 1 and so forth. Rows were filtered using a list of 25 RNAi defense-related genes to produce a submatrix with bulk data for 23 of those genes (Mp6g20400 and Mp5g06390 were not included in the original Seurat). Values are then changed to z-scores using R’s *scale* function. Finally, all cells with z-scores corresponding to TPM values lower than 1 were changed to NA so that the corresponding cells in the final heatmap are eventually rendered white for easy identification. The heatmap plot was then produced using *R*’s *pheatmap* package. All genes with TPM values lower than 1 in all cell clusters were removed from the final plot. The above described strategy was also used for analysis related to PPR and NLR genes. Dot plots were produced using the matrix of Marchantia z-score values explained above. For each cell type and each member of the different defense mechanisms, the percentage of expressed genes (number of genes with non-NA values relative to the total number of RNAi genes detected in all three cell types) was computed, as well as the corresponding average z-scores. These values were then plotted using *ggplot2*.

For cell type specific transcriptomes and their response to induction of P1/Hc-Pro viral silencing suppressor, multiplexed stranded libraries were produced from 500 pgrs of total RNA (RIN>8) using the SMART-Seq® v4 Ultra® Low Input RNA Kit (Takara) and sequenced in an Illumina HiSeqX device. A total of 17-27 million pairs (150 bp reads) were obtained after preprocessing steps with *Trim Gallore!* and *SortMeRNA*, as described above. Quantification was performed using *Salmon* v1.5.1, with same parameters as stated above, and the Arabidopsis transcriptome file AtRTD2_19April2016.fa (https://ics.hutton.ac.uk/atRTD/) modified by adding the transcript sequences from TuMV P1/Hc-Pro and mScarlet-I.The **R** program *tximport* v1.14.2 was then used to aggregate *Salmon*’s transcript expression estimates at gene level. Outliers were detected and removed using https://www.graphpad.com/quickcalcs/Grubbs1.cfm based on the level of induction of scarlet (control samples) and P1/HcPro to select the most similar three biological replicates from the original four per sample type for further analyses. A preliminary filter was applied to exclude lowly expressed genes, removing those with fewer than 10 total counts across all analyzed samples. This reduced the number of genes from 33,684 to 26,386.

Genes from each defense pathway were considered expressed when at least two of the three biological replicates showed expression levels ≥1 TPM, and the mean expression across those same replicates was also ≥1 TPM. Average TPM values were then calculated for each set of three biological replicates. For genes deemed absent in a given cell type, the corresponding average TPM values were set to zero, even if they exceeded 1. In addition, all average TPM values below 1 were set to zero. Z-scores were subsequently calculated, and dot plots were generated as described above for *Marchantia*.

Differential expression analysis in Arabidopsis samples was carried out using DESeq2’s^51^ *results* function with the filtered and normalized object as input and *alpha* parameter set to *0*.*05*.

Functional information was added to the results using the R package *biomaRt*.

Gene Ontology analysis were performed using the ShinyGO v0.75 (http://bioinformatics.sdstate.edu/go/) tool with default parameters (FDR<0.05) and selecting the option of showing 30 top pathways to show.

### Phenotyping

Seeds from primary transformants for the lines expressing P1/Hc-Pro under HTR5, CAB3, Sultr2;2 and SUC promoters were selected based on the expression of mcherry in the seed coat. Plants were grown randomized in LD conditions and 22º C. Two-week-old plants were sorted into 4 categories according to the phenotype penetrance inferred by the severity of developmental abnormalities.

## Data availability

Raw sequencing data have been deposited in the European Nucleotide Archive (ENA) under accession number PRJEB113077.

## Acknowledgments

We would like to thank to Miguel Ángel Blazquez and Salomé Prat for critical reading of the manuscript. We are grateful to Bruno Pok Man Ngou for assistance in obtaining PRR annotation in Arabidopsis and Marchantia. We are also grateful to Bozeng Tand and Wenbo Ma for sharing their NLR annotation in Arabidopsis. The Green gate module containing the constitutive Arabidopsis HTR5 gene promoter (At4g40400) was kindly provided by Mathieu Ingouff. Plant codon optimized SlmScarlet was kindly provided by Idan Efroni. Work at the MoRE Lab was funded by RTI2018-097262-B-100 and PID2024-163067NB-100 (funded by MCIN/AEI/ 10.13039/501100011033 and by “ESF Investing in your future”). L.V.-M. was supported by BES-2016-076986 and B.M. was supported by PREP2024-002308 (Contracts PREP2024-PID) (both funded by MCIN/AEI/ 10.13039/501100011033 and by “ESF Investing in your future”). B.M. T.J.G. was recipient of a Postdoctoral Fellowship from CRAG through the “Severo Ochoa Program for Centres of Excellence in R&D” 2016-2019 (SEV-2015-0533).

## Figures and tables

**Table 1: DNA constructs used in this study**.

**Table 2: List of primers used in this study**.

## References

1 Kappagantu, M., Collum, T. D., Dardick, C. & Culver, J. N. Viral Hacks of the Plant Vasculature: The Role of Phloem Alterations in Systemic Virus Infection. Annu Rev Virol 7, 351–370, doi:10.1146/annurev-virology-010320-072410 (2020).

2 Hipper, C., Brault, V., Ziegler-Graff, V. & Revers, F. Viral and cellular factors involved in Phloem transport of plant viruses. Front Plant Sci 4, 154, doi:10.3389/fpls.2013.00154 (2013).

3 Ros-Moner, E. et al. Conservation of molecular responses upon viral infection in the non-vascular plant Marchantia polymorpha. Nat Commun 15, 8326, doi:10.1038/s41467-024-52610-0 (2024).

4 Gimenez-Ibanez, S., Zamarreno, A. M., Garcia-Mina, J. M. & Solano, R. An Evolutionarily Ancient Immune System Governs the Interactions between Pseudomonas syringae and an Early-Diverging Land Plant Lineage. Curr Biol 29, 2270–2281 e2274, doi:10.1016/j.cub.2019.05.079 (2019).

5 Carella, P. et al. Conserved Biochemical Defenses Underpin Host Responses to Oomycete Infection in an Early-Divergent Land Plant Lineage. Curr Biol 29, 2282–2294 e2285, doi:10.1016/j.cub.2019.05.078 (2019).

6 Redkar, A. et al. Marchantia polymorpha model reveals conserved infection mechanisms in the vascular wilt fungal pathogen Fusarium oxysporum. New Phytol 234, 227–241, doi:10.1111/nph.17909 (2022).

7 Ngou, B. P. M., Jones, J. D. G. & Ding, P. Plant immune networks. Trends Plant Sci 27, 255–273, doi:10.1016/j.tplants.2021.08.012 (2022).

8 Silvestri, A., Bansal, C. & Rubio-Somoza, I. After silencing suppression: miRNA targets strike back. Trends Plant Sci 29, 1266–1276, doi:10.1016/j.tplants.2024.05.001 (2024).

9 Ngou, B. P. M., Kadota, Y. & Shirasu, K. Plant cell surface receptors. Plant J 125, e70800, doi:10.1111/tpj.70800 (2026).

10 Niehl, A., Wyrsch, I., Boller, T. & Heinlein, M. Double-stranded RNAs induce a pattern-triggered immune signaling pathway in plants. New Phytol 211, 1008–1019, doi:10.1111/nph.13944 (2016).

11 Ngou, B. P. M., Wyler, M., Schmid, M. W., Kadota, Y. & Shirasu, K. Evolutionary trajectory of pattern recognition receptors in plants. Nat Commun 15, 308, doi:10.1038/s41467-023-44408-3 (2024).

12 Barragan, A. C. & Weigel, D. Plant NLR diversity: the known unknowns of pan-NLRomes. Plant Cell 33, 814–831, doi:10.1093/plcell/koaa002 (2021).

13 Lopez-Gomollon, S. & Baulcombe, D. C. Roles of RNA silencing in viral and non-viral plant immunity and in the crosstalk between disease resistance systems. Nat Rev Mol Cell Biol 23, 645–662, doi:10.1038/s41580-022-00496-5 (2022).

14 Belanger, S., Zhan, J. & Meyers, B. C. Phylogenetic analyses of seven protein families refine the evolution of small RNA pathways in green plants. Plant Physiol 192, 1183–1203, doi:10.1093/plphys/kiad141 (2023).

15 Dombey, R. et al. Atypical epigenetic and small RNA control of degenerated transposons and their fragments in clonally reproducing Spirodela polyrhiza. Genome Res 35, 522–544, doi:10.1101/gr.279532.124 (2025).

16 Ernst, E. et al. Duckweed genomes and epigenomes underlie triploid hybridization and clonal reproduction. Curr Biol 35, 1828–1847 e1829, doi:10.1016/j.cub.2025.03.013 (2025).

17 Li, S. X., Liu, Y., Zhang, Y. M., Chen, J. Q. & Shao, Z. Q. Convergent reduction of immune receptor repertoires during plant adaptation to diverse special lifestyles and habitats. Nat Plants, doi:10.1038/s41477-024-01901-x (2025).

18 Ngou, B. P. M., Heal, R., Wyler, M., Schmid, M. W. & Jones, J. D. G. Concerted expansion and contraction of immune receptor gene repertoires in plant genomes. Nat Plants 8, 1146–1152, doi:10.1038/s41477-022-01260-5 (2022).

19 Wang, L. et al. The maturation and aging trajectory of Marchantia polymorpha at single-cell resolution. Dev Cell 58, 1429–1444 e1426, doi:10.1016/j.devcel.2023.05.014 (2023).

20 Chia, K. S. et al. The N-terminal domains of NLR immune receptors exhibit structural and functional similarities across divergent plant lineages. Plant Cell 36, 2491–2511, doi:10.1093/plcell/koae113 (2024).

21 Aubry, S., Smith-Unna, R. D., Boursnell, C. M., Kopriva, S. & Hibberd, J. M. Transcript residency on ribosomes reveals a key role for the Arabidopsis thaliana bundle sheath in sulfur and glucosinolate metabolism. Plant Journal 78, 659–673, doi:10.1111/tpj.12502 (2014).

22 Takada, S. & Goto, K. TERMINAL FLOWER2, an homolog of HETEROCHROMATIN PROTEIN1, counteracts the activation of FLOWERING LOCUS T by CONSTANS in the vascular tissues of leaves to regulate flowering time. Plant Cell 15, 2856–2865, doi:10.1105/tpc.016345 (2003).

23 Li, D. et al. Arabidopsis Class II TCP Transcription Factors Integrate with the FT-FD Module to Control Flowering. Plant Physiol 181, 97–111, doi:10.1104/pp.19.00252 (2019).

24 Tian, S., Luo, X., Cui, B. & He, Y. Auto-downregulation of the florigen FT production prevents precocious flowering in plants. bioRxiv, 2025.2004.2024.650415, doi:10.1101/2025.04.24.650415 (2025).

25 Olas, J. J. et al. Nitrate acts at the Arabidopsis thaliana shoot apical meristem to regulate flowering time. New Phytol 223, 814–827, doi:10.1111/nph.15812 (2019).

26 Wu, G. & Poethig, R. S. Temporal regulation of shoot development in Arabidopsis thaliana by miR156 and its target SPL3. Development 133, 3539–3547, doi:10.1242/dev.02521 (2006).

27 Xu, T. et al. Expressional and regulatory characterization of Arabidopsis RNA-dependent RNA polymerase 1. Planta 237, 1561–1569, doi:10.1007/s00425-013-1863-7 (2013).

28 Zhou, M. et al. The CLASSY family controls tissue-specific DNA methylation patterns in Arabidopsis. Nat Commun 13, 244, doi:10.1038/s41467-021-27690-x (2022).

29 Tang, B., Feng, L., Hulin, M. T., Ding, P. & Ma, W. Cell-type-specific responses to fungal infection in plants revealed by single-cell transcriptomics. Cell Host Microbe 31, 1732–1747 e1735, doi:10.1016/j.chom.2023.08.019 (2023).

30 Bedoya, L. C., Martinez, F., Orzaez, D. & Daros, J. A. Visual tracking of plant virus infection and movement using a reporter MYB transcription factor that activates anthocyanin biosynthesis. Plant Physiol 158, 1130–1138, doi:10.1104/pp.111.192922 (2012).

31 Pollari, M., De, S., Wang, A. & Makinen, K. The potyviral silencing suppressor HCPro recruits and employs host ARGONAUTE1 in pro-viral functions. PLoS Pathog 16, e1008965, doi:10.1371/journal.ppat.1008965 (2020).

32 Sanobar, N. et al. Investigating the Viral Suppressor HC-Pro Inhibiting Small RNA Methylation through Functional Comparison of HEN1 in Angiosperm and Bryophyte. Viruses 13, doi:10.3390/v13091837 (2021).

33 Valli, A. A., Gallo, A., Rodamilans, B., Lopez-Moya, J. J. & Garcia, J. A. The HCPro from the Potyviridae family: an enviable multitasking Helper Component that every virus would like to have. Mol Plant Pathol 19, 744–763, doi:10.1111/mpp.12553 (2018).

34 Schurholz, A. K. et al. A Comprehensive Toolkit for Inducible, Cell Type-Specific Gene Expression in Arabidopsis. Plant Physiol 178, 40–53, doi:10.1104/pp.18.00463 (2018).

35 Xue, H. C. et al. A unified cell atlas of vascular plants reveals cell-type foundational genes and accelerates gene discovery. Cell 188, 6370–6390 e6329, doi:10.1016/j.cell.2025.07.036 (2025).

36 El Mahboubi, K. et al. Plant-fungi interactions in Marchantia polymorpha are associated with horizontal gene transfer and terpene metabolism. Proc Natl Acad Sci U S A 123, e2532723123, doi:10.1073/pnas.2532723123 (2026).

37 Espinosa-Cores, L., Michavila, S., Gonzalez-Zuloaga, M., Solano, R. & Gimenez-Ibanez, S. MpNPR modulates lineage-specific oil body development and defence against gastropod herbivory in <em>Marchantia polymorpha</em>. bioRxiv, 2025.2011.2017.688000, doi:10.1101/2025.11.17.688000 (2025).

38 Nobori, T. et al. A rare PRIMER cell state in plant immunity. Nature 638, 197–205, doi:10.1038/s41586-024-08383-z (2025).

39 Zhu, J. et al. Single-cell profiling of Arabidopsis leaves to Pseudomonas syringae infection. Cell Rep 42, 112676, doi:10.1016/j.celrep.2023.112676 (2023).

40 Beck, M. et al. Expression patterns of flagellin sensing 2 map to bacterial entry sites in plant shoots and roots. J Exp Bot 65, 6487–6498, doi:10.1093/jxb/eru366 (2014).

41 Yotsui, I. et al. LysM-mediated signaling in Marchantia polymorpha highlights the conservation of pattern-triggered immunity in land plants. Curr Biol 33, 3732–3746 e3738, doi:10.1016/j.cub.2023.07.068 (2023).

42 Lampropoulos, A. et al. GreenGate---a novel, versatile, and efficient cloning system for plant transgenesis. PLoS One 8, e83043, doi:10.1371/journal.pone.0083043 (2013).

43 Ranjan, A., Fiene, G., Fackendahl, P. & Hoecker, U. The Arabidopsis repressor of light signaling SPA1 acts in the phloem to regulate seedling de-etiolation, leaf expansion and flowering time. Development 138, 1851–1862, doi:10.1242/dev.061036 (2011).

44 Takahashi, H. et al. The roles of three functional sulphate transporters involved in uptake and translocation of sulphate in Arabidopsis thaliana. Plant J 23, 171–182, doi:10.1046/j.1365-313x.2000.00768.x (2000).

45 Truernit, E. & Sauer, N. The promoter of the Arabidopsis thaliana SUC2 sucrose-H+ symporter gene directs expression of beta-glucuronidase to the phloem: evidence for phloem loading and unloading by SUC2. Planta 196, 564–570, doi:10.1007/BF00203657 (1995).

46 Todesco, M., Rubio-Somoza, I., Paz-Ares, J. & Weigel, D. A collection of target mimics for comprehensive analysis of microRNA function in Arabidopsis thaliana. PLoS Genet 6, e1001031, doi:10.1371/journal.pgen.1001031 (2010).

47 Weigel D & Glazebrook J. Arabidopsis: A Laboratory Manual. Cold Spring Harbor Laboratory Press, Cold Spring Harbor, N.Y., ©2002 (2002).

48 Blazquez, M. A., Soowal, L. N., Lee, I. & Weigel, D. LEAFY expression and flower initiation in Arabidopsis. Development 124, 3835–3844, doi:10.1242/dev.124.19.3835 (1997).

49 Patro, R., Duggal, G., Love, M. I., Irizarry, R. A. & Kingsford, C. Salmon provides fast and bias-aware quantification of transcript expression. Nat Methods 14, 417-+, doi:10.1038/nmeth.4197 (2017).

50 Soneson, C., Love, M. I. & Robinson, M. D. Differential analyses for RNA-seq: transcript-level estimates improve gene-level inferences. F1000Res 4, 1521, doi:10.12688/f1000research.7563.2 (2015).

51 Love, M. I., Huber, W. & Anders, S. Moderated estimation of fold change and dispersion for RNA-seq data with DESeq2. Genome Biol 15, 550, doi:10.1186/s13059-014-0550-8 (2014).

