## Supplementary Table 5 for "Developmental diversification re-patterns basal antiviral immunity across plant cell types"

| Construct IDs | Construct modules | Purpose |
|---|---|---|
| pTJ1 | CAB3p:SV40NLS-dTomato-RBCSt, AT2S3p:mCherry-MASt in pGGZ003 | Mesophyll cells labeling |
| pTJ41 | CAB3p:SV40NLS-eGFP- RBCSt, AT2S3p:mCherry-MASt in pGGZ003 | Mesophyll cells labeling |
| pTJ44 | Sultr2;2p:SV40NLS-eGFP- RBCSt, AT2S3p:mCherry-MASt in pGGZ003 | Bundle sheath cells labeling |
| pTJ43 | SUC2p:SV40NLS-eGFP- RBCSt, AT2S3p:mCherry-MASt in pGGZ003 | Companion cells labeling |
| pAP12 | AtRDR6p:GUS-RDR6 3'UTR, AT2S3p:mCherry-MASt in pGGZ003 | AtRDR6 transcriptional reporter |
| pN108 | HTR5p:SV40NLS-COmScarlet-RBCSt in pGGM000 | sRNA activity reporter |
| pLV59 | HTR5p:AmiRCOmScarlet-RBCSt, NOSp:NeoR in pGGN000 | sRNA activity reporter |
| pN167 | pN108+pLV59 in pGGZ003 | sRNA activity reporter |
| pKU2 | HTR5p:P1/Hc-Pro-UBIQ10t, AT2S3p:mCherry-MASt in pGGZ003 | Constitutive P1/Hc-Pro expression |
| pLV188 | CAB3p:P1/Hc-Pro-UBIQ10t, AT2S3p:mCherry-MASt in pGGZ003 | Mesophyll P1/Hc-Pro expression |
| pLV190 | Sultr2;2p:P1/Hc-Pro-UBIQ10t, AT2S3p:mCherry-MASt in pGGZ003 | Bundle sheath P1/Hc-Pro expression |
| pLV189 | SUC2p:P1/Hc-Pro-UBIQ10t, AT2S3p:mCherry-MASt in pGGZ003 | Companion cells P1/Hc-Pro expression |
| pLV90 | CAB3p:GR-LhG4-RBCSt in pGGM000 | Dex induction in Mesophyll |
| pLV146 | Sultr2;2p: GR-LhG4-RBCSt in pGGM000 | Dex induction in Bundle sheath |
| pLV93 | SUC2p: GR-LhG4-RBCSt in pGGM000 | Dex induction in Companion cells |
| pLV100 | 6xOPp:P1/Hc-Pro-UBIQ10t, UBIQ10p :HygR-OCSt in pGGN000 | Induction of P1/Hc-Pro |
| pLV108 | pLV90+pLV100 in pGGZ003 | P1/Hc-Pro induction in Mesophyll |
| pLV148 | pLV146+pLV100 in pGGZ003 | P1/Hc-Pro induction in Bundle sheath |
| pLV110 | pLV93+pLV100 in pGGZ003 | P1/Hc-Pro induction in Companion cells |
| pLV126 | 6xOPp:SV40NLS-COmScarlet, UBIQ10p :HygR-OCSt in pGGN000 | Induction of COmScarlet |
| pLV124 | pLV90+pLV126 in pGGZ003 | NLS-COmScarlet induction in Mesophyll |
| pLV147 | pLV146+pLV126 in pGGZ003 | NLS-COmScarlet induction in Bundle sheath |
| pLV149 | pLV93+pLV126 in pGGZ003 | NLS-COmScarlet induction in Companion cells |

**Supplementary Table 5. Constructs used in this study.**
