## Supplementary figures and images for "Developmental diversification re-patterns basal antiviral immunity across plant cell types"

### Supplementary Figure 1

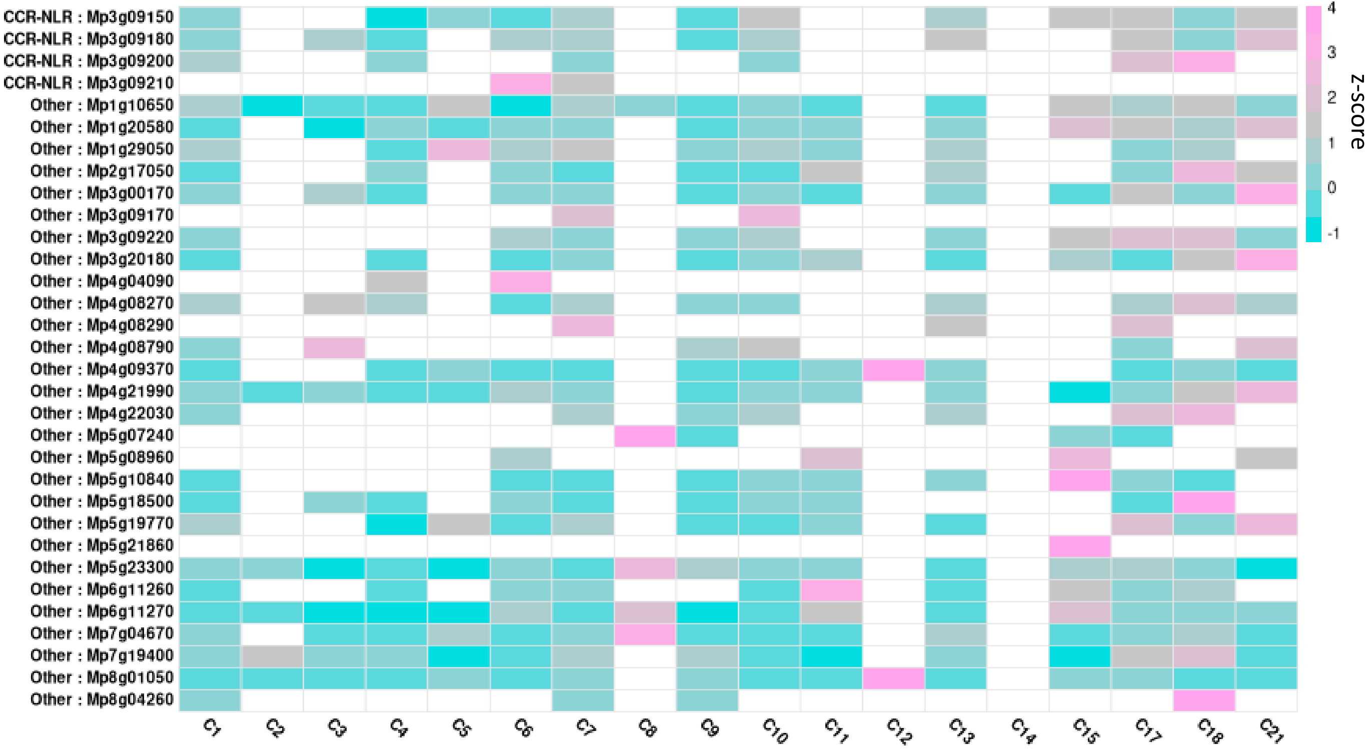

### Supplementary Figure 2

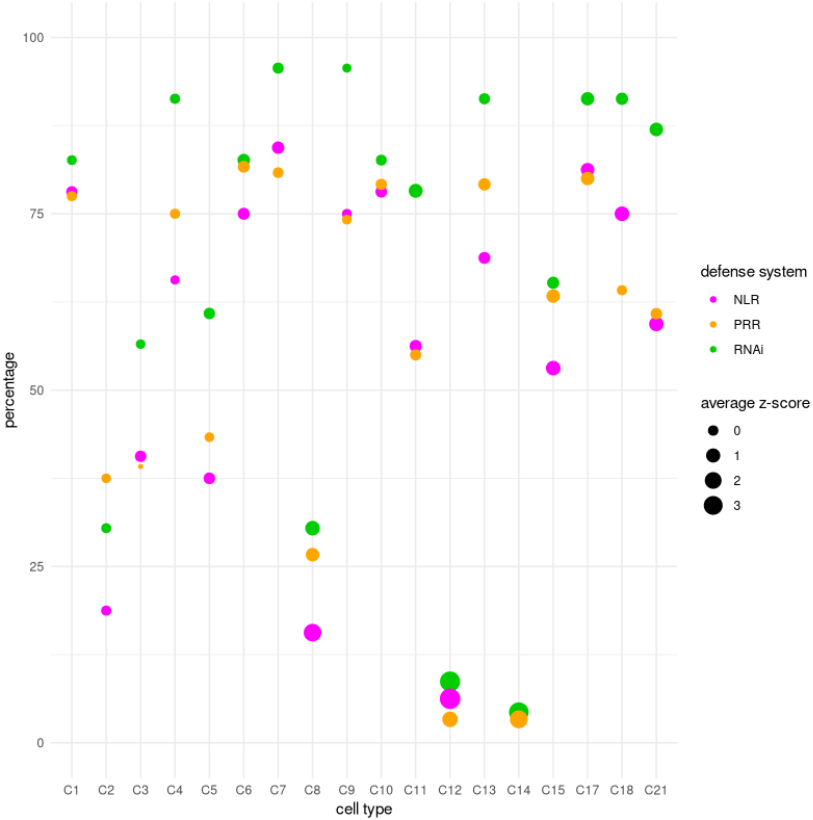

### Supplementary Figure 3

#### Mp RLPs

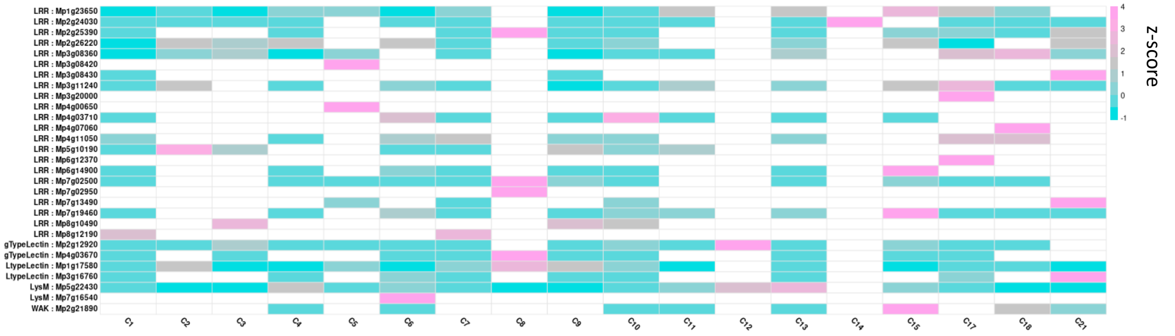


#### Mp RLKs

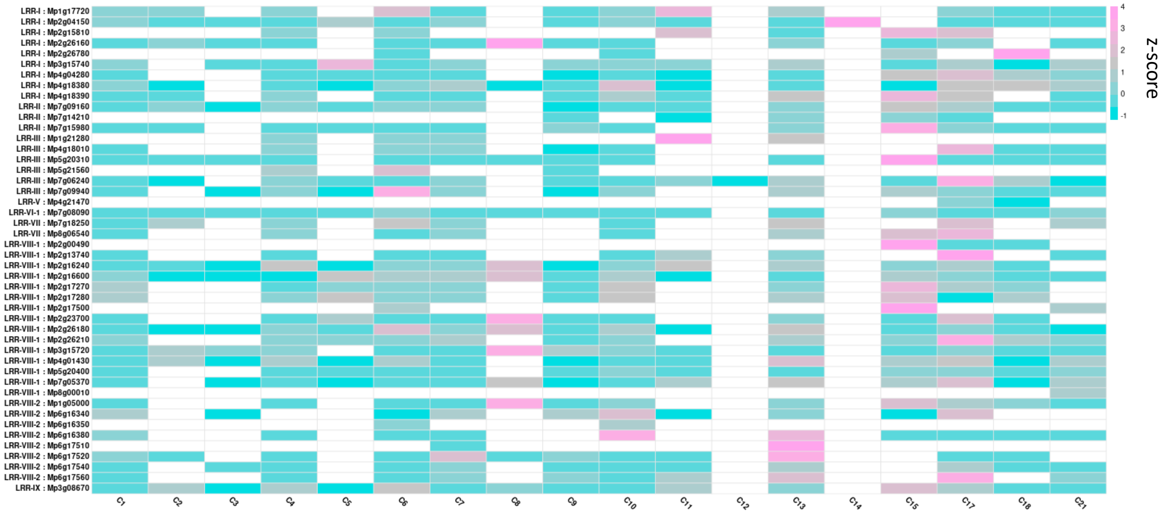


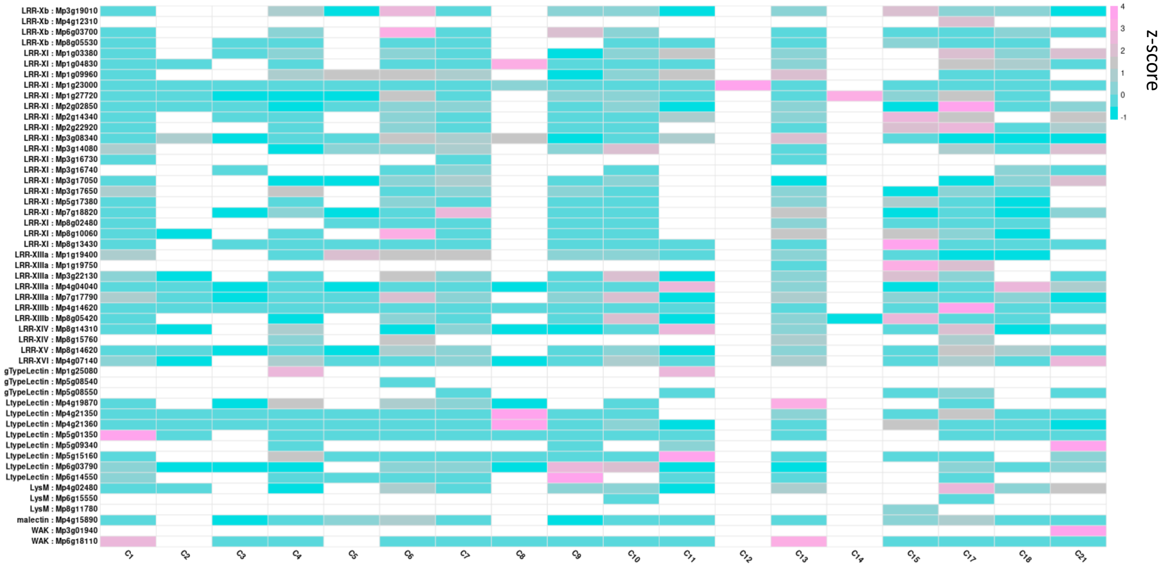

### Supplementary Figure 4

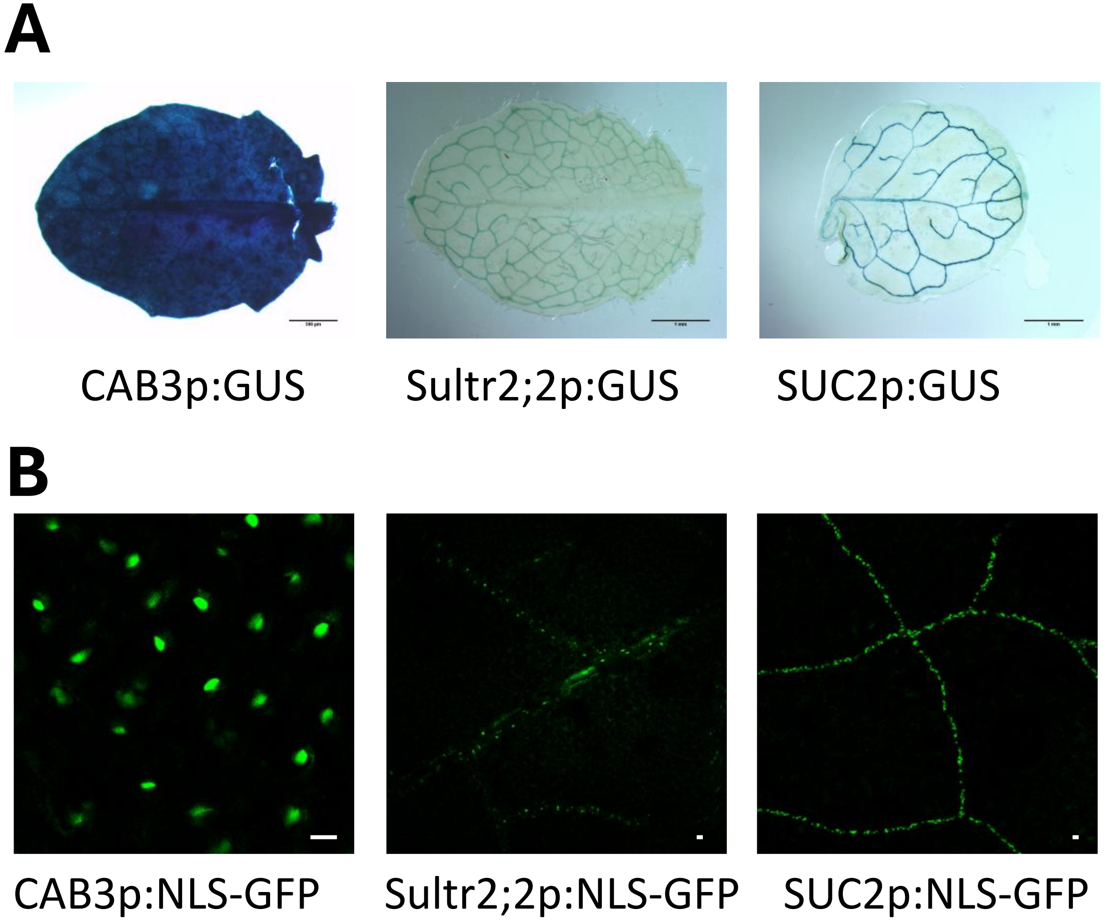

### Supplementary Figure 5

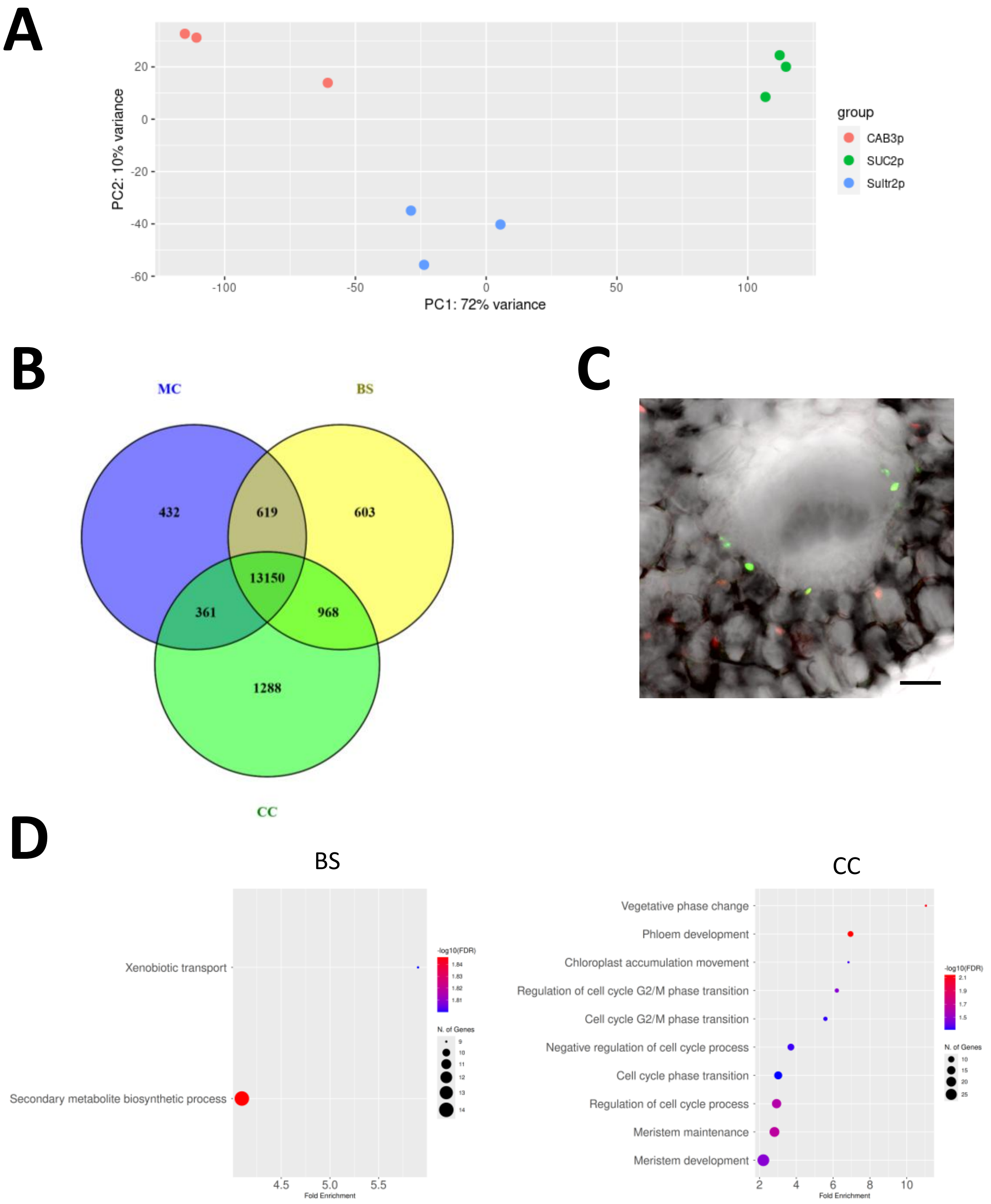

### Supplementary Figure 6

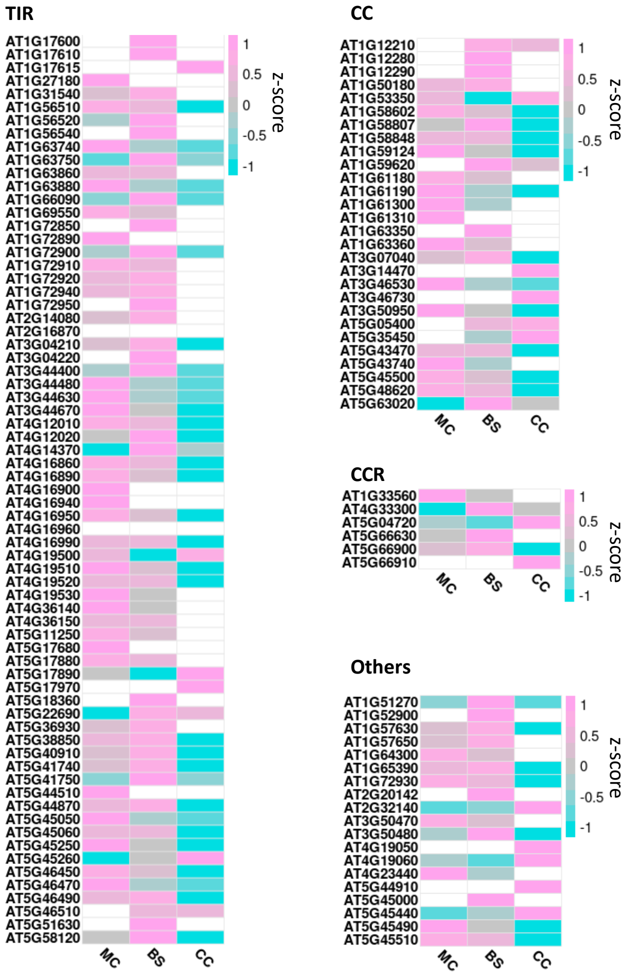

### Supplementary Figure 7

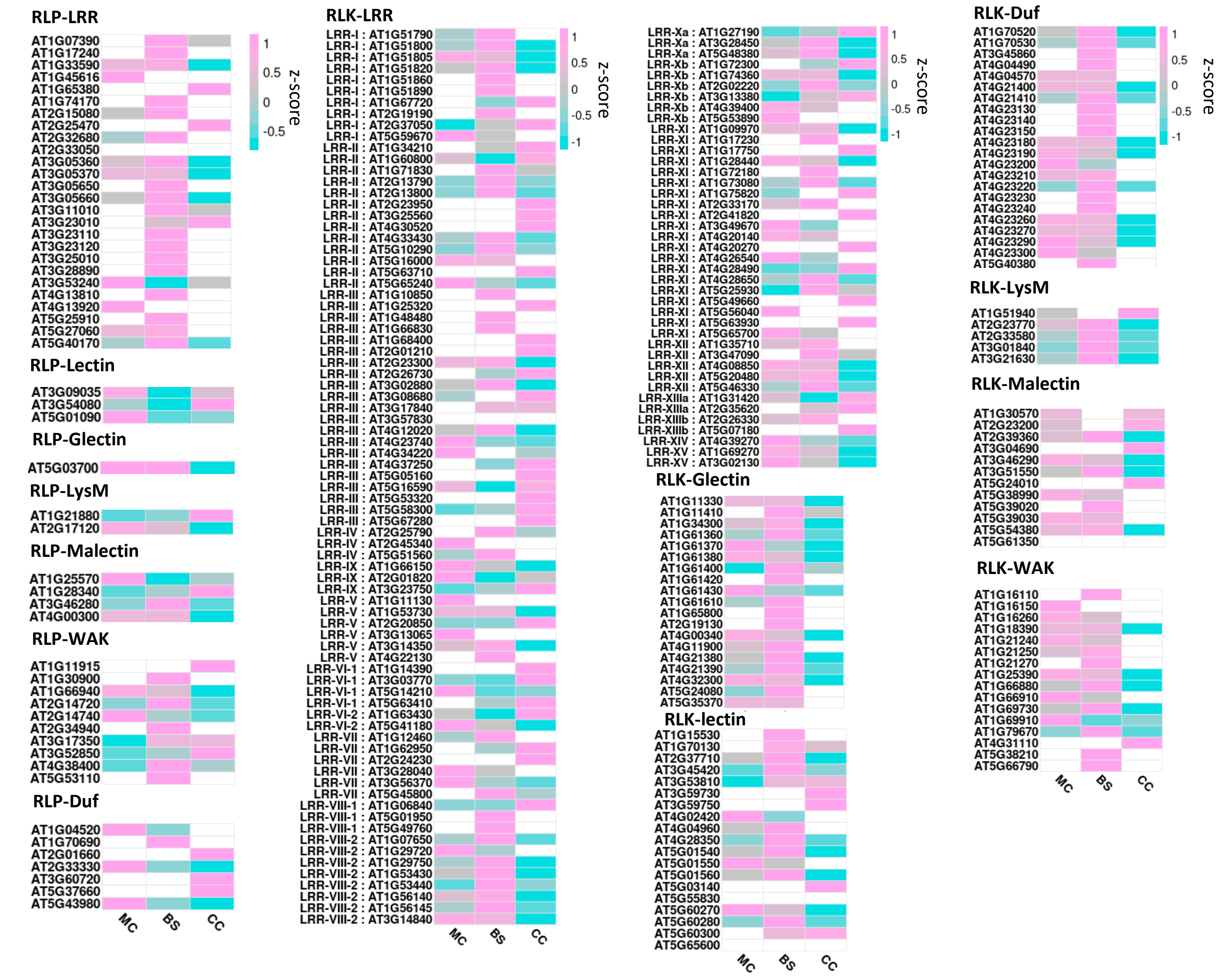

### Supplementary Figure 8

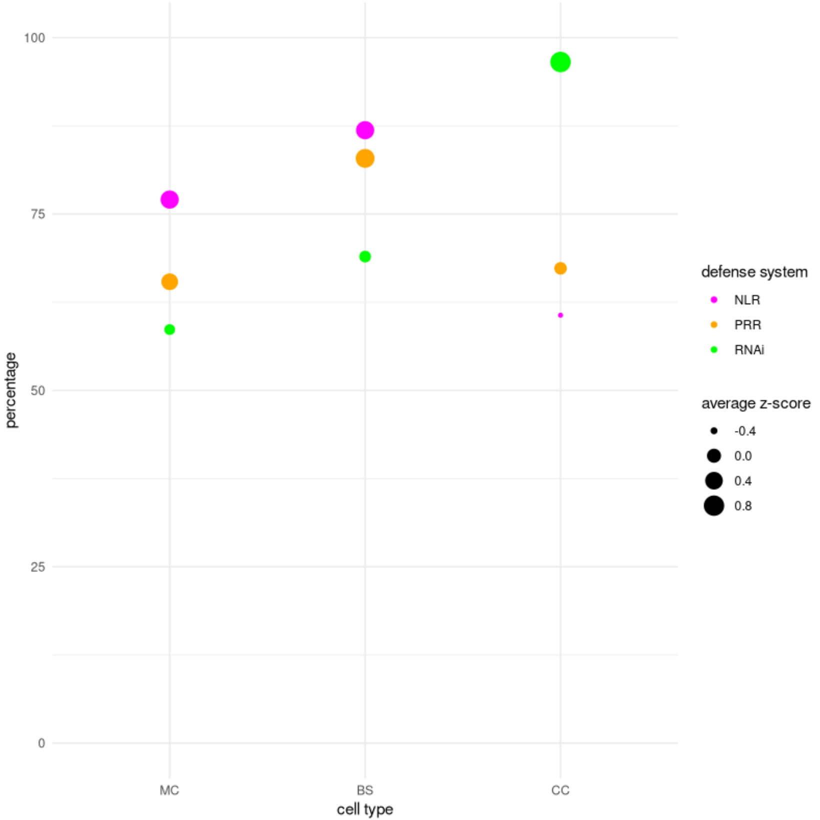

### Supplementary Figure 9

A

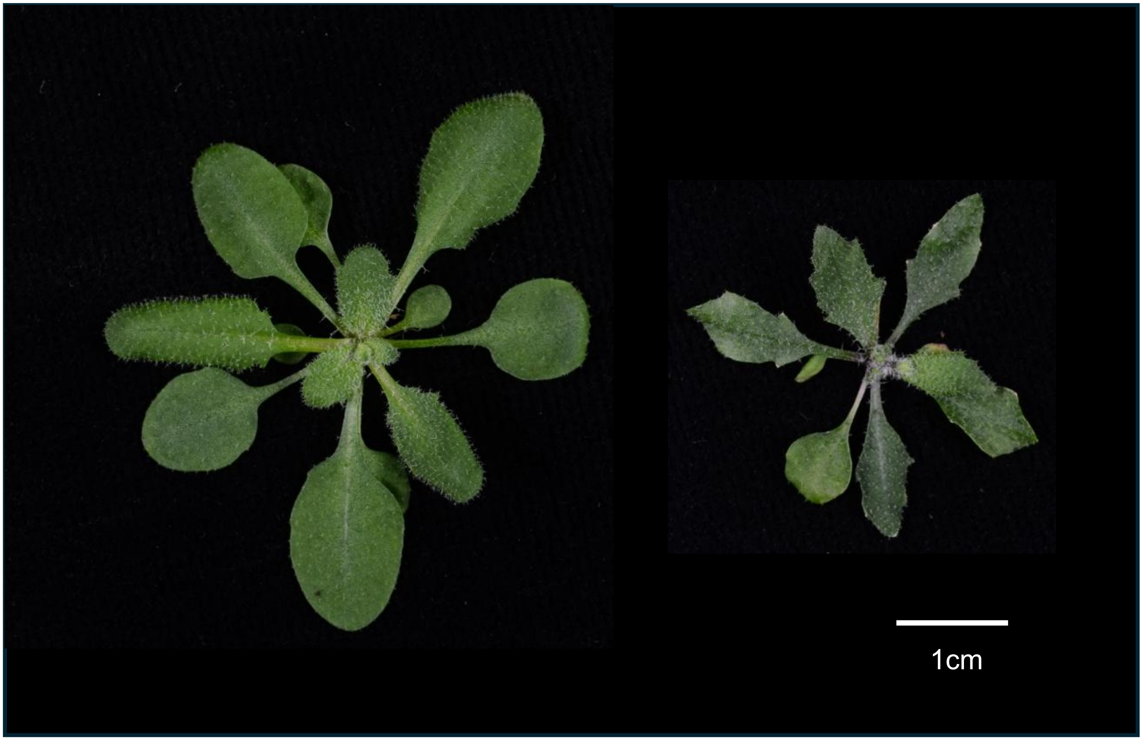


B

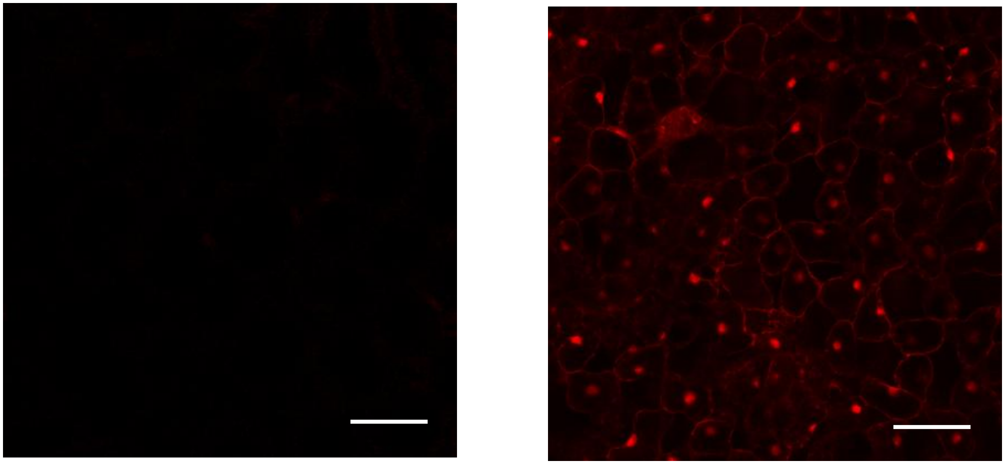

### Supplementary Figure 10

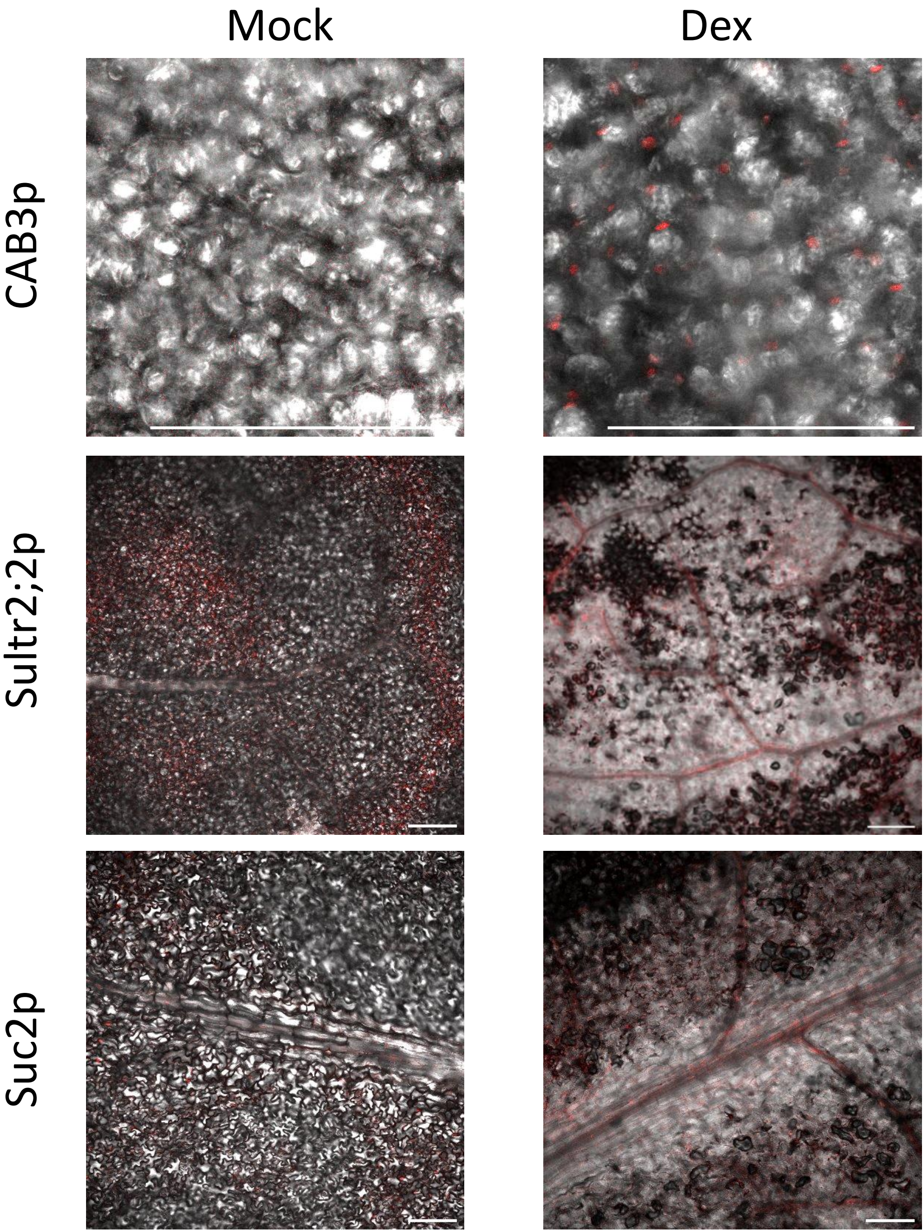
