## Supplementary Table 4 for "Developmental diversification re-patterns basal antiviral immunity across plant cell types"

| Primer IDs | Common name | Sequence (5'-3') | Purpose |
|---|---|---|---|
| M-136 | CAB3p.A.s | GGTCTCaACCTGATTATGTTCTTTTCAAA | Cloning CAB3p |
| M-137 | CAB3p.B.as | GGTCTCtTGTTTGAAACTTTTTGTGTTTT | Cloning CAB3p |
| M-142 | CAB3pmut.s | GTGGTCTAGAAATGCTTTGG | Eco31I site mutation CAB3p |
| M-143 | CAB3pmut.as | CATTTCTAGACCACATGTTGC | Eco31I site mutation CAB3p |
| M-379 | Sultr2;2p.A.s | aacaGGTCTCaacctaaccattattttatagc aattttacataaatatatttagatattg | Cloning Sultr2;2p |
| M-380 | Sultr2;2p.B.as | aacaGGTCTCttgttTCAGCTCTCTCTCT AGATATATATTAACTTTTTT | Cloning Sultr2;2p |
| M-398 | Sultr2;2.m1.s | attatgagagtccaaacgtttcaaatttgtc | Eco31I site mutation Sultr2;2p |
| M-399 | Sultr2;2.m1.as | tttggactctcataatataagaaaccgatcga | Eco31I site mutation Sultr2;2p |
| M-400 | Sultr2;2.m2.s | ggaaaaagaggcccaaaaagaaaagaac | Eco31I site mutation Sultr2;2p |
| M-401 | Sultr2;2.m2.as | tgggcctctttttccttatctttttatatggaag | Eco31I site mutation Sultr2;2p |
| M-71 | P1/Hc-Pro.B.s | GGTCTCaAACAATGGCAGCAGTTACA | Cloning P1/Hc-Pro |
| M-72 | P1/Hc-Pro.C.as | GGTCTCaAGCCCTAGAGTGCGTAATCTG | Cloning P1/Hc-Pro |
| M-244 | SlmScarlet-iCO.C.s | aacaGGTCTCaggctcaacaATGGTGAGC AAGGGCG | SlmScarlet-iCO cloning |
| M-245 | SlmScarlet-iCO.D.as | aacaGGTCTCtctgaTCACGAACCTCTAG ATCCGG | SlmScarlet-iCO cloning |
| M-199 | amiR.B.s | aaaGGTCTCaAACAacaaacacacgctcgg | AmiR cloning |
| M-200 | amiR.C.as | aaaGGTCTCaAGCCcatggcgatgccttaaataa ag | AmiR cloning |
| M-246 | amiRScarletCO.1.I.s | gaTCTGCACGGGCTTCTTGGCCActc tcttttgtattcca | AmiR construction |

| | | | |
|---|---|---|---|
| M-247 | amiRScarletCO..1II.as | agTGGCCAAGAAGCCCGTGCAGAtcaaagagaatcaatga | AmiR construction |
| M-248 | amiRScarletCO.1.III.s | agTGACCAAGAAGCCGGTGCAGTtcacaggtcgtgatatg | AmiR construction |
| M-249 | amiRScarletCO.1.IV.as | gaACTGCACCGGCTTCTTGGTCActacatatatattccta | AmiR construction |
| M-1061 | RDR6p.A.s | caGGTCTCaacctaaatgaacttcaaacactatacttgaaaaag | AtRDR6p cloning |
| M-1062 | RDR6p.B.as | caGGTCTCttgttttctcctgaaaaagaaacatactaaacag | AtRDR6p cloning |
| M-1080 | RDR6-3UTR.C.s | caGGTCTCaggctaaggttatgtttaatgccgtaagg | AtRDR6-3'UTR cloning |
| M-1081 | RDR6-3UTR.F.as | caGGTCTCatagtcgttttctcaccaaaacgaggaag | AtRDR6-3'UTR cloning |

**Supplementary Table 4. Primers used in this study**
